# Phosphorylation-Dependent Activation of the bHLH Transcription Factor ICE1/SCRM Promotes Polarization of the Arabidopsis Zygote

**DOI:** 10.1101/2024.07.03.601926

**Authors:** Houming Chen, Feng Xiong, Torren Bischoff, Kai Wang, Yingjing Miao, Daniel Slane, Rebecca Schwab, Thomas Laux, Martin Bayer

## Abstract

Asymmetric cell divisions are a key mechanism for breaking symmetry and orchestrating different cell identities in multicellular organisms. In *Arabidopsis thaliana*, as in most flowering plants, the first zygotic cell division is asymmetric, giving rise to the embryo proper and an extraembryonic suspensor.

Zygotic polarization and differential cell identities in the daughter cells are controlled by the ERECTA-YODA pathway, a prototype receptor kinase-MAP kinase signaling pathway. This pathway also controls asymmetric cell divisions in the epidermis during stomatal development. In this context, the bHLH transcription factor ICE1/SCRM is a direct target of MPK3/6, and phosphorylation negatively controls SCRM activity by targeting the protein for proteasomal degradation. This raises the question if this regulatory module is also involved in the asymmetric division of the zygote.

Our results show that SCRM has a critical function in zygote polarization and acts in parallel with the known MPK3/6 target WRKY2 in activating the homeobox transcription factor gene *WOX8*. Our work further demonstrates that SCRM activity in the early embryo is positively controlled by MPK3/6-mediated phosphorylation. Therefore, the mode of MAP kinase regulation of the same target protein fundamentally differs between the embryo and the epidermis, shedding light on cell type-specific, differential gene regulation by common signaling pathways.

## Introduction

Multicellularity enables organisms to develop specialized cell and tissue types with dedicated functions. Asymmetric cell divisions are key to breaking symmetry and initiating distinct developmental programs in daughter cells (Sunchu & Cabernard, 2020; Guo & Dong, 2022; Sugioka, 2022; Bogaert *et al*., 2023).

In many angiosperms, embryogenesis is initiated by an asymmetric division of the zygote, accompanied by the establishment of a primary apical-basal axis (Lindsey & Topping, 1993). In Arabidopsis, this asymmetric division results in a smaller apical cell with an embryonic fate and a larger basal cell that contributes primarily to the extraembryonic suspensor (Mansfield & Briarty, 1991). Polarity establishment in the Arabidopsis zygote is controlled by the evolutionarily conserved ERECTA-YODA mitogen-activated protein (MAP) kinase pathway (Lukowitz *et al*., 2004; Chen *et al*., 2021). Currently, known members of this pathway comprise receptor kinases of the ERECTA (ER) family (ERf) (Torii *et al*., 1996; Shpak *et al*., 2005; Wang *et al*., 2021) as well as two members of the membrane-associated BRASSINOSTEROID SIGNALING KINASE (BSK) family, BSK1 and BSK2 (Neu *et al*., 2019; Wang *et al*., 2021) that transduce signaling to the MAP kinase cascade. The sperm cell-derived BSK protein SHORT SUPENSOR (SSP/BSK12), which represents a naturally occurring constitutively active variant of BSK protein, can activate MAP kinase signaling independent of ERf function (Bayer *et al*., 2009; Neu *et al*., 2019; Wang *et al*., 2021). The MAP kinase cascade includes the MAP3K YODA (YDA) (Lukowitz *et al*., 2004), MAP KINASE KINASE 4 (MKK4) and MKK5, as well as the MAP KINASE 3 (MPK3) and MPK6 (Wang *et al*., 2007; Zhang, M *et al*., 2017). MPK3/6 activate the downstream transcription factor (TF) WRKY2 in a phosphorylation-dependent manner to regulate the early zygote and suspensor-specific gene *WUS-RELATED HOMEOBOX 8* (*WOX8*) (Ueda, M. *et al*., 2011; Ueda *et al*., 2017). Defects in zygote elongation and polar division plane positioning are morphological hallmarks of reduced YDA activation and are easily detectable in *er* single*, er erl2* double, *ssp* single, *bsk1 bsk2* double and *wrky2* single mutants (Bayer *et al*., 2009; Ueda *et al*., 2017; Neu *et al*., 2019; Wang *et al*., 2021).

While our understanding of YDA signaling in the early zygote has increased significantly in the last years, most of the conceptual knowledge about this pathway has been gained by studying its function in stomata development, a process that is similarly initiated by asymmetric cell divisions (Guo *et al*., 2021; Herrmann & Torii, 2021; Smit & Bergmann, 2023). Interestingly, while cell lineage initiation in stomata development and zygote polarization seem to rely by and large on the same upstream signal transduction components, known downstream TF targets seem to be different.

During the first steps of stomata development, ERf- and BSK1/2-dependent activation of the YDA-MKK4/5-MPK3/6 cascade leads to phosphorylation of the basic helix-loop-helix (bHLH) transcription factor SPEECHLESS (SPCH), marking it for ubiquitination and degradation (MacAlister, C. A. *et al*., 2007). SPCH promotes the asymmetric entry division that forms a smaller meristemoid mother cell and a larger stomata lineage ground cell (MacAlister, C. A. *et al*., 2007; Wang *et al*., 2007; Lampard, Gregory R *et al*., 2008). This first step is reminiscent of YDA function in the early embryo. MUTE and FAMA, two close homologs of SPCH, fulfil consecutive functions in the development of the meristemoid to guard cells. MUTE promotes the differentiation of the meristemoid to a guard mother cell, and is, as in the case of SPCH, negatively controlled by ERECTA-YODA signaling (MacAlister, Cora A *et al*., 2007; Pillitteri *et al*., 2008; Qi *et al*., 2017). A single round of cell division in the guard mother cell and final differentiation of the pair of guard cells is controlled by FAMA (Ohashi-Ito & Bergmann, 2006). Here, a MAP kinase cascade including YDA, MKK7/9, and MPK3/6 positively regulates activity (Lampard *et al*., 2009). However, MUTE and FAMA are not a substrate for MPK6-mediated phosphorylation as they lack a MAP kinase target domain (MPKTD).

SCREAM (SCRM), also named INDUCER OF CBF EXPRESSION 1 (ICE1), a broadly expressed bHLH protein, has been reported as a central regulator in both freezing tolerance and stomata development (Chinnusamy *et al*., 2003; Kanaoka *et al*., 2008). Transcriptional activity of SCRM is essential for induction of *C-REPEAT BINDING FACTOR* (*CBF*) and further cold-responsive genes in cold stress (Chinnusamy *et al*., 2003; Lee *et al*., 2005; Kim *et al*., 2015). Further studies have identified SCRM as a phosphorylation substrate of the MKK4/5-MPK3/6 cascade and MPK3/6-directed phosphorylation on SCRM protein negatively regulates tolerance of freezing *in planta* (Li *et al*., 2017; Zhao *et al*., 2017). In stomata development, SCRM and its close paralogue SCRM2 form heterodimers with SPCH, MUTE and FAMA to orchestrate stomata initiation, commitment, and terminal differentiation, respectively (Kanaoka *et al*., 2008). Loss of SCRM leads to stomata defects and *scrm scrm2* double mutants phenocopy *spch* mutants that lack mature guard cells on the cotyledon surface (Kanaoka *et al*., 2008). Recent research has shown that SCRM functions as a scaffold to bridge MPK3/6 and SPCH via a bipartite motif, the MAPK docking site and the KRAAM sequence (Putarjunan *et al*., 2019). In a previously isolated gain-of-function mutant *scrm-D,* a single amino acid substitution in the KRAAM motif (R236H) disrupts normal interaction between MPK3/6 and SCRM and blocks degradation of SCRM (Putarjunan *et al*., 2019). Consequently, the *scrm-D* mutation leads to increased activity of the SPCH/SCRM-D heterodimer and a strong gain-of-function phenotype with an epidermis almost entirely consisting of stomata.

Given the prominent function of the ERECTA-YODA pathway in controlling asymmetric divisions in stomata and embryo development, we asked whether SCRM would have a similar function in the embryo as in the epidermis. Here, we show that SCRM is also a target of the embryonic YDA pathway and plays a crucial role in zygote polarization and establishment of different cell identities in the early embryo. However, our results suggest that, in contrast to its stomatal function, SCRM activity is positively controlled by MPK-directed phosphorylation. Furthermore, genetic evidence indicates that SCRM functions in parallel with the previously described transcription factor WRKY2 to activate *WOX8* in the zygote. Our work not only refines our knowledge of the embryonic YDA pathway and the activation of early patterning genes in plants, but also highlights the context-specific control of transcription factor activity. The same MPK3/6-dependent phosphorylation of SCRM can lead to a differential tissue-specific activation or inactivation of target genes.

## Methods and materials

### Plant Materials and Growth Conditions

Arabidopsis strains used in this research are all in Columbia-0 (Col-0) background and were grown in a walk-in plant growth chamber with a 16 h: 8 h, light: dark photoperiod at 23 ℃. The mutants in this study were described previously: *scrm* (*ice1-2*, SALK_003155), *scrm-D*(R236>H)*, wrky2-1*(SALK_020399), *scrm2-1* (SAIL_808_B10), and *yda-11* (SALKseq_078777) (Kanaoka *et al*., 2008; Pal *et al*., 2017; Ueda *et al*., 2017; Ohta *et al*., 2018; Wang *et al*., 2021). The gWOX8△-YFP marker line and ICE1-GFP 6A were published previously (Ueda, Minako *et al*., 2011; Li *et al*., 2017). Genotyping primers and details are listed in the Supplementary Table1.

### Molecular Cloning and Generation of Transgenic Plants

In-fusion cloning by the In-Fusion HD cloning Kit (TAKARA) was used for plasmid construction. The *YPet* coding sequence (CDS) following a *NOS* terminator was cloned into SalI and KpnI restriction sites in a *pBay-Hyg* vector, a modified pCambia vectors detailed in Wang *et al*., 2021, yielding *YPet-pBay-Hyg*. An *SCRM* genomic fragment containing 2589 bp sequence upstream of the start codon and 1835 bp genomic region without a stop codon was in-frame introduced between BamHI and SalI into *YPet-pBay-Hyg* to generate *SCRM-YPet*. For *YPet-SCRM*, a 5134 bp upstream region of *SCRM* was first cloned into BcuI/BamHI sites of *YPet-pBay-Hyg* backbone, then the *NOS* terminator fragment was excised by ApaI/KpnI digest and replaced by a 1838 bp *SCRM* genomic sequence with stop codon and 611 bp downstream sequence. *SCRM* genomic fragments were amplified from genomic DNA with Q5^®^ High-fidelity polymerase (NEB). *YPet-SCRM-D, YPet-SCRM-6A* were produced by site-directed mutagenesis PCR using *YPet-SCRM* as a template. In-Fusion cloning and mutagenesis primers were included in Supplementary Table2.

The binary vectors were transformed into *Agrobacterium tumefaciens* GV3101 by electroporation for the floral dip method in Arabidopsis transformation (Clough & Bent, 1998). The positive transformants were screened on 1/2 Murashige and Skoog media containing 1%(w/v) sucrose, 0.8%(w/v) agar and 20 mg/L hygromycin.

### CRISPR/Cas9-Generated Plant Mutants

The CRISPR system in this research was established according to former methods. Plant codon-optimized *pcoCas9* sequence (Li *et al*., 2013) combined with *rbcS* terminator driven by the egg-cell specific promoter *pEC1.2-EC1.1* (Wang *et al*., 2015) was cloned into *pBay-Bar* backbone and the endosperm-specific promoter *pAT2S3* connected with *mCherry* sequence (Gao *et al*., 2016) was subsequently introduced to generate CRISPR destination vector *pBay-Bar-CRmCherry*. To produce the sgRNA cassettes, a pair of 20 bp target sequences were picked according to CRISPR-P 2.0 (Liu *et al*., 2017) and introduced into shuttle-in vectors pEF004/005 via mutagenesis PCR as described (Wu *et al*., 2018). The two sgRNA cassettes were subcloned into *pBay-Bar-CRmCherry* between XmaJI and MluI by In-Fusion assembly. The destination constructs were then transformed into Col-0 wildtype by the floral dip method (Clough & Bent, 1998). Positive T1 plants were selected by phosphinothricin resistance on 1/2 MS media containing 50 mg/L phosphinothricin and Cas9-free T2 or T3 mutants were identified by absence of mCherry marker. To generate *scrm-cr* and *ssp-6*, sgRNA cassettes were designed to remove genomic sequence (Fig. S1b) and the genotyping primers are detailed in the Supplementary Table1.

### Microscopy Imaging

Differential interference contrast (DIC) imaging with a Zeiss Axio Imager was applied to measure the lengths of apical/basal cells at the 1-cell stage and suspensors at transition stage from ovule samples, which were dissected and cleared overnight in Hoyer’s solution as described in Bayer *et al*., 2009. Length measurements were conducted by AxioVision 4 and ZEN 2.0 blue edition software.

SCRM-YPet embryonic expression and gWOX8△-YFP signals were detected by a Zeiss LSM 780 NLO microscope (Laser with 514nm wavelength for excitation, detectors record emission for wavelength between 517nm and 570nm). Immature seeds were immersed in desalt water and nuclear signal intensity in zygotes was measured by ZEN 2.0 blue edition software.

For propidium iodide (PI) staining, cotyledons from 5-day-old seedlings were dissected and submerged in 10 μg/ml PI (Sigma–Aldrich) for 10 min. Cotyledon samples were rinsed 3 times with water and then transferred to slides with abaxial side on the top. PI signal detection (571-656nm) was under excitation of 561nm wavelength laser and measured by Fiji 2.0.0 and ZEN 2.0 blue edition software.

### Split-Luciferase Complementation Assay

A split-luciferase complementation assay was conducted as described (Zhou *et al*., 2018). Full-length CDS of *SCRM* and *WRKY2* were cloned into *pCAMBIA1300-Nluc* and *pCAMBIA1300-Cluc* respectively. Meanwhile *SPCH-pCAMBIA1300-Cluc* was generated as a positive control and empty vector *pCAMBIA1300-Cluc* functioned as a negative control. These plasmids were produced between digestion sites KpnI and SalI by In-Fusion assembly and the cloning primers were collected in the Supplementary Table2. These constructs were transferred into *Agrobacterium tumefaciens* GV3101 and positive clones were cultured for 16 h and suspended in induction buffer (10 mM MES pH5.7, 10 mM MgCl_2_ and 150 mM acetosyringone) with OD600 at 0.5. Agrobacterial suspension was infiltrated into leaves of 4-week-old *Nicotiana benthamiana* from abaxial side and tobaccos were kept in the plant chamber for 48 h. Then leaf discs (0.6 cm diameter) from plants were poked out and immersed into luciferin substrate (1 mM luciferin aqueous solution with 0.02% Silwet L-77) to detect relative luminescence by Infinite F200 Fluorescence Microplate Reader (Tecan). For western blotting, SCRM-Nluc was detected by Anti-luciferase antibody (Sigma, L0159) and CLuc constructs were immunoblotted by Anti-CLuc antibody (Sigma, L2164).

### Dual-Luciferase Assay

The dual-luciferase assay was carried out in essence as published before (Hellens *et al*., 2005) with some modifications. 2511 bp length of *WOX8* promoter was cloned into reporter plasmid *pGREENii-0800LUC* with BcuI and NcoI by In-Fusion reaction to produce *pWOX8-LUC*. For the *pWOX8cisA-LUC* reporter plasmid, *35s* minimal promoter was first introduced into *pGREENii-0800LUC between* BcuI and NcoI, and then 416 bp *WOX8* cisA element was inserted in the upstream using BamHI and BcuI. Effector constructs were generated by inserting CDS of SCRM and bHLH35 after 2 x 35S promoters in pJIT60 vectors. Reporter and effector plasmids were co-transformed into 125 µl protoplast cell suspension with the standard protoplast transfection protocol (Mehlhorn *et al*., 2018) in Transformation Facility of the Center for Plant Molecular Biology (ZMBP), University of Tübingen. The luciferase activity was detected using Dual-Luciferase^®^ Reporter Assay System (Promega, E1910) by Infinite F200 Fluorescence Microplate Reader (Tecan).

### Statistical Analysis of Quantitative Data

One-way ANOVA analysis with tukey test and student’s t test and were used for statistical analysis. Diagrams and statistical analyses were made by Prism 9.3 (https://www.graphpad.com/scientific-software/prism/) and BoxPlotR (Spitzer *et al*., 2014).

## Results

### SCRM is a critical factor in zygote polarization

The asymmetric division of the zygote relies on a similar MPK-dependent pathway as the asymmetric entry division of the stomata lineage and *SCRM* transcripts can be detected in the zygote (Fig. S1a) (Zhao *et al*., 2019). Therefore, we hypothesized that the initiation of embryogenesis relies on a similar function of SCRM.

To determine whether SCRM is a downstream target of the YDA pathway in embryogenesis, we first performed phenotypic analyses of loss-of-function alleles. At the 1-cell stage of the embryo, the combined length of the apical and basal cell represents the final length of the zygote, since cell division occurs after the elongation phase (Mansfield & Briarty, 1991; Kang *et al*., 2023). The ratio of apical to basal cell size can therefore be used as a quantitative readout of cell polarization as it directly reflects the asymmetric, apical positioning of the division plane in the zygote (Ueda *et al*., 2017; Wang *et al*., 2021).

For these measurements, we used a previously published *scrm* T-DNA null allele (Kanaoka *et al*., 2008) as well as a novel CRISPR *scrm-cr* mutant allele (Fig. S1b). Compared to wildtype, *scrm* mutants exhibited reduced zygote length (Fig. 1a, b, Fig. S1c) as well as impaired asymmetry (Fig. 1c). Since *yda* homozygous plants are sterile and do not produce embryos, we used a CRISPR/Cas9-mediated deletion mutation in the *SSP* gene, termed *ssp-6* (Fig. S1b), for direct comparison of the phenotype.

**Figure 1:**
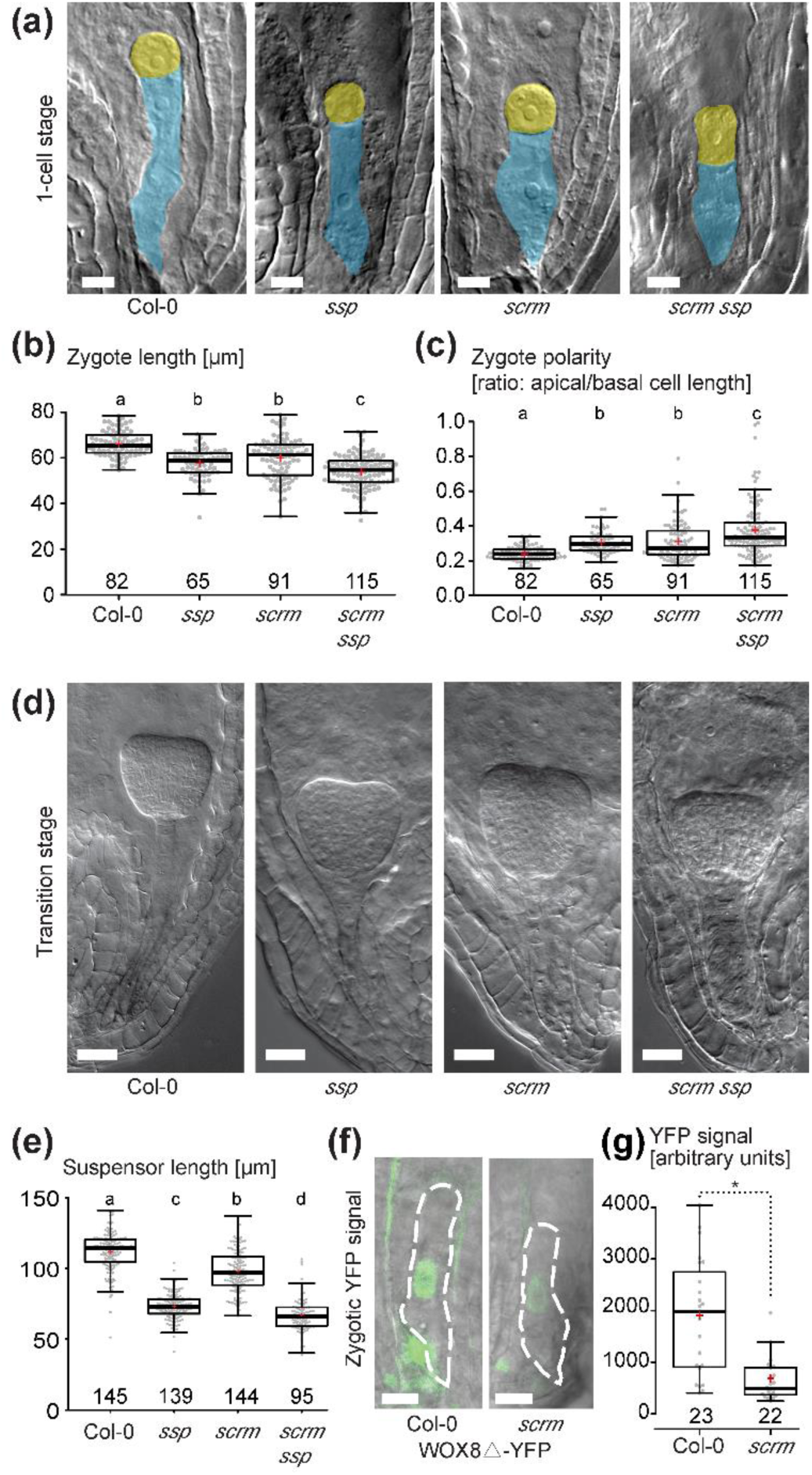
**SCRM is required for zygotic polarization and elongation.** (a) DIC images of one cell-stage embryos (apical cells false-colored in yellow and basal cells colored in blue). Genotypes are as indicated below figure panel. Scale bars, 10 µm. (b, c) Boxplot diagram of zygote lengths (sum of apical and basal cell lengths at 1-cell stage in b) and zygotic polarity (ratios of apical and basal cell lengths in c). (d) DIC images of transition stage embryos. Genotypes are indicated below figure panels. Scale bar, 20 µm. (e) Boxplot diagram of suspensor lengths at transition stage. (f) Fluorescence microscopy images of nuclear gWOX8Δ-YFP signal in zygotes of wildtype and *scrm* whole-mount ovules. Scale bar, 10 µm. (g) Boxplot diagram of gWOX8Δ-YFP signal intensity in wildtype and *scrm*. In (b, c, e, g), the sample size is given above the x axis. Center lines show the medians; box limits indicate the 25th and 75th percentiles; whiskers extend 1.5 times the interquartile range from the 25th and 75th percentiles; red crosses represent sample means; data points are plotted as gray dots. Letters above boxes refer to individual groups in a one-way ANOVA with a post hoc Tukey test (p < 0.05). In g, statistically significant differences in Student t-test (p<0.05) are indicated by an asterisk.

Stronger phenotypes were observed in the *scrm ssp-6* double mutant than in the *scrm* or *ssp* single mutant (Fig. 1a-e). Since loss of *SSP* does not lead to a complete loss of YDA-dependent signaling (Bayer *et al*., 2009; Wang *et al*., 2021), additional loss of *SCRM* would further impair the pathway and could explain this additive effect. At the transition stage, a similar trend was observed for suspensor lengths (Fig. 1d, e). Impaired zygote elongation and polarization and compromised suspensor development are features common to mutants of the embryonic ERECTA-YODA signaling pathway and suggest that SCRM is an integral part of this pathway.

The homeodomain transcription factor gene *WOX8* is a target of the embryonic YODA pathway, since its expression in the zygote and the early embryo is controlled by MPK6-dependent phosphorylation of the transcription factor WRKY2 (Ueda *et al*., 2017). To test if SCRM can influence *WOX8* expression, we analyzed the activity of a transcriptional *WOX8* reporter (*gWOX8Δ-YFP* (Ueda, M. *et al*., 2011) in the *scrm* mutant and wildtype background. The YFP signal in the zygote was significantly reduced in *scrm* compared to wildtype (Fig. 1f, g), suggesting that *WOX8* expression partially depends on *SCRM* function.

Taken together, these data suggest that SCRM is required for strong activation of *WOX8* expression and zygote polarization.

### SCRM functions in parallel with WRKY2

Measurements of *WOX8* reporter gene expression in *scrm* mutant zygotes showed reduced but still detectable activity (Fig. 1f, g). Furthermore, embryonic defects in *scrm* single mutants are less severe than observed in *yda* mutants (Fig. 1d, e; Fig. S3a, b). Therefore, other factors must be working in parallel to SCRM in controlling *WOX8* activity in the zygote.

The closely related gene *SCRM2*, which has largely overlapping functions with *SCRM* in stomatal development, does not appear to be expressed in embryos based on the available transcriptome data (Fig. S1a (Zhao *et al*., 2019)), and no zygotic or suspensor defects were observed in *scrm2* (Fig. S2a-c). Furthermore, no enhancement of the *scrm* mutant phenotype can be observed in *scrm/SCRM scrm2* double mutants (Fig. S2a-c). Since *scrm scrm2* homozygous double mutants are sterile, the analysis was performed using a segregating population of embryos from heterozygous *scrm/SCRM* parents considering that *scrm* mutants are zygotic recessive.

The transcription factor WRKY2 has previously been described as target of the embryonic ERECTA-YODA pathway (Ueda *et al*., 2017), and the reduced *WOX8* expression in *scrm* mutants is reminiscent of WRKY2 function.

To understand the genetic relationship between *SCRM* and *WRKY2*, we analyzed *scrm* and *wrky2-1* single mutants and *scrm wrky2-1* double mutants. Simultaneous loss of *SCRM* and *WRKY2* strongly enhanced the embryonic phenotypes of *scrm* or *wrky2* single mutants with extremely short zygotes and almost complete loss of zygote polarization, reminiscent of homozygous *yda* embryos (Fig. 2a, b, c). At later developmental stages, aberrant embryo development with a barely recognizable suspensor became apparent (Fig. S3a, b), similar to homozygous *yda* embryos (Lukowitz *et al*., 2004). The effect on *gWOX8Δ-YFP* reporter gene expression in the *scrm wrky2* double mutant followed this trend, showing a further reduction in signal intensity compared to the single mutants and wild-type (Fig. 2d).

**Figure 2:**
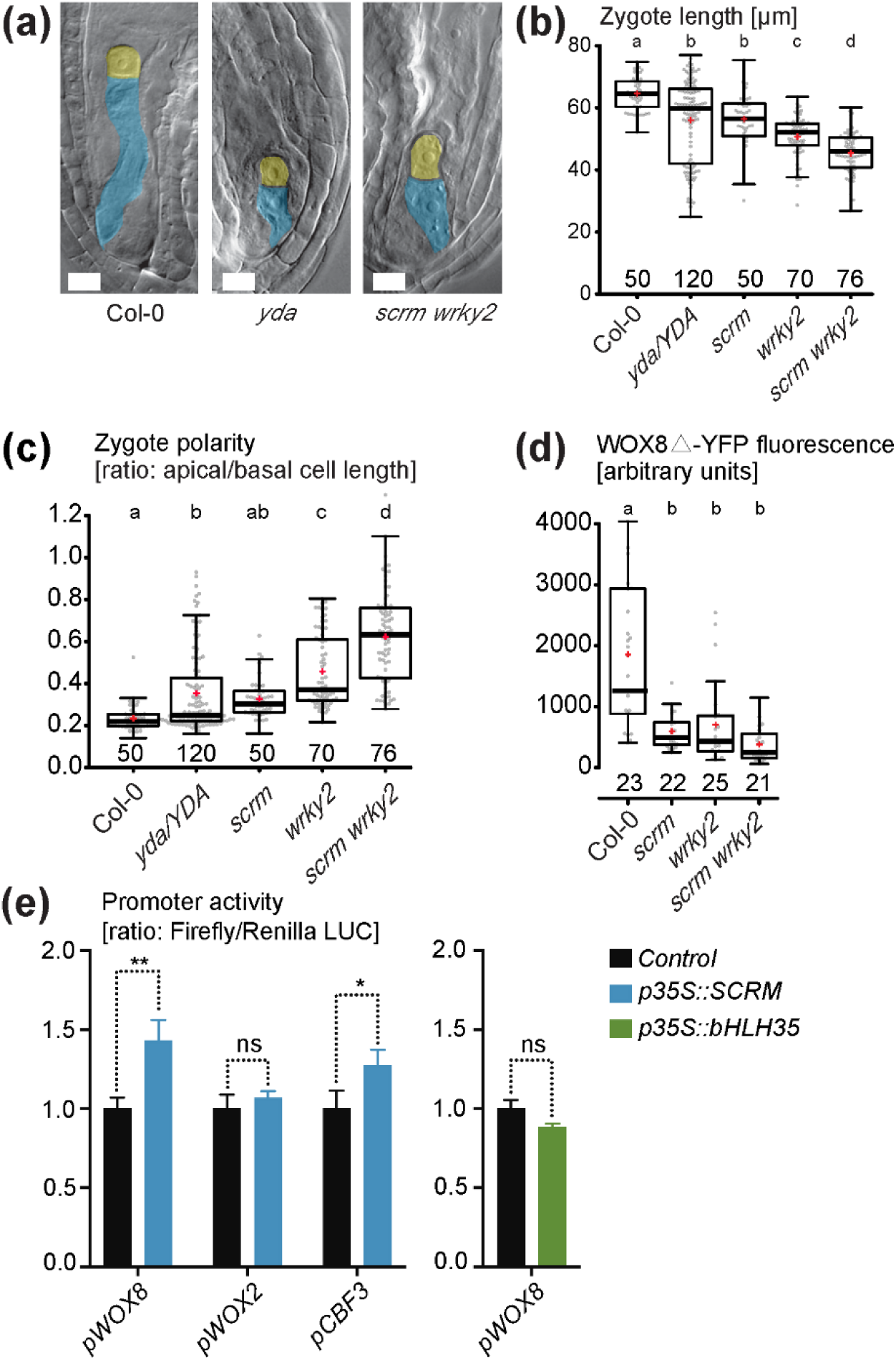
**SCRM and WRKY2 work synergistically in zygote polarization.** (a) DIC images of one cell-stage embryos (apical cells false-colored in yellow and basal cells colored in blue). Genotypes are as indicated below figure panel. Scale bars, 10 µm. (b, c) Boxplot diagram of zygote lengths (sum of apical and basal cell lengths at 1-cell stage in b) and zygotic polarity (ratios of apical and basal cell lengths in c). (d) Boxplot diagram of *gWOX8-YFP* fluorescent signal intensity in zygotes. In (b-d), the sample size is given above the x axis. Center lines show the medians; box limits indicate the 25th and 75th percentiles; whiskers extend 1.5 times the interquartile range from the 25th and 75th percentiles; red crosses represent sample means; data points are plotted as gray dots. Letters above boxes refer to individual groups in a one-way ANOVA with a post hoc Tukey test (p < 0.05). (e) Dual luciferase activity assay in protoplasts of suspension cell cultures. Data is illustrated as means ± SD of three independent experimental replicates, asterisk indicates statistical differences in Student’s t test (p<0.05); ns (p>0.05). Color of the bar indicates which transcription factor gene was co-expressed as indicated in the legend (Control = empty vector). The promoter sequences used to control firefly luciferase expression are given below each set of bar graphs.

To investigate the possibility that these proteins could act as heterodimers, we tested for protein interaction using a split-luciferase assay. We were unable to detect any interaction between SCRM and WRKY2 (Fig. S3c, d), whereas the previously published interaction of the SCRM-SPCH heterodimer as a control was readily detectable. This suggests that SCRM and WRKY2 do not act as a heterodimeric complex in *WOX8* activation. The additive effects of *scrm* and *wrky2* mutants in the early embryo could be a further indication that these two transcription factors act independently on the same target gene. We used a 2.5 kb promoter fragment of *WOX8* in a dual luciferase assay in Arabidopsis suspension cell protoplasts to test for activation of *WOX8* by SCRM (Fig. 2e). As suggested by the reduced *WOX8* reporter gene expression in *scrm* mutants, we detected increased expression of the *pWOX8::LUC* construct in the presence of SCRM, whereas there was no increase in expression of *pWOX8::LUC* by the closely related transcription factor bHLH35 or the *pWOX2::LUC* control construct. As a positive control, we included a previously described direct target of SCRM in cold stress response as a chimeric *pCBF3::LUC* reporter, which was activated in this assay to the same extend as *pWOX8::LUC* (Fig. 2e). These results suggest that SCRM can specifically activate *WOX8* expression.

In a previous study, cis-regulatory elements for *WOX8* activation by WRKY2 were mapped to a region termed *cisB* in the second intron of *WOX8* (Ueda et al., 2011). In the same study, an additional region upstream of the start codon (*cisA*) was identified as a site of *WOX8* activation (Ueda, Minako *et al*., 2011). Since we were able to detect activation of *LUC* expression by SCRM with a construct containing the *cisA* region, we tested this fragment in a dual luciferase assay. LUC activity was strongly increased by SCRM when this element was fused to a 35S minimal promoter (Fig. S3e). Taken together, these results suggest that *WOX8* activation by SCRM is direct but might be independent of and additive to WRKY2 function.

### Embryonic functions of SCRM rely on phosphorylation

In the asymmetric entry division of the stomatal lineage, SCRM activity is negatively influenced by the ERECTA-YODA pathway as is the case during cold stress response (Li *et al*., 2017; Zhao *et al*., 2017). However, the phenotype of *scrm* loss-of-function mutants is similar to that of loss-of-function mutants of the embryonic YDA pathway, suggesting a positive control of SCRM activity by the ERECTA-YODA MAPK pathway, opposite to the wiring in epidermal patterning. This raises the question of whether SCRM is also a direct target of MPK3/6-mediated phosphorylation in the embryonic YDA pathway or whether SCRM activity is controlled in a different manner. To address this question, we analyzed the embryonic phenotype of the stomatal gain-of-function mutant *scrm-D* (Fig. 3a-d) (Kanaoka *et al*., 2008). *scrm-D* is a missense mutation that disrupts MPK3/6 interaction and MPK3/6-dependent phosphorylation of the SCRM-D protein. The single amino acid substitution (R236H) in the KRAAM motif results in ectopic SCRM protein accumulation and strong gain-of-function phenotypes during stomata development (Fig. S4a) (Kanaoka *et al*., 2008; Putarjunan *et al*., 2019). Interestingly, reduced zygote lengths and shorter suspensors were detected in *scrm-D* embryos at the one-cell stage and transition stage (Fig. 3a-d), reminiscent of *scrm* loss-of-function alleles. Loss of zygote polarization was more pronounced in *scrm-D* than in *scrm* (Fig. 3b), suggesting a dominant negative effect. A newly generated *YPet-scrm-D* transgenic line also essentially mimicked the phenotypes of *scrm-D*, showing more stomata in cotyledons (Fig. S4a, b) and reduced zygote elongation and polarization (Fig. S4c, d).

**Fig 3.**
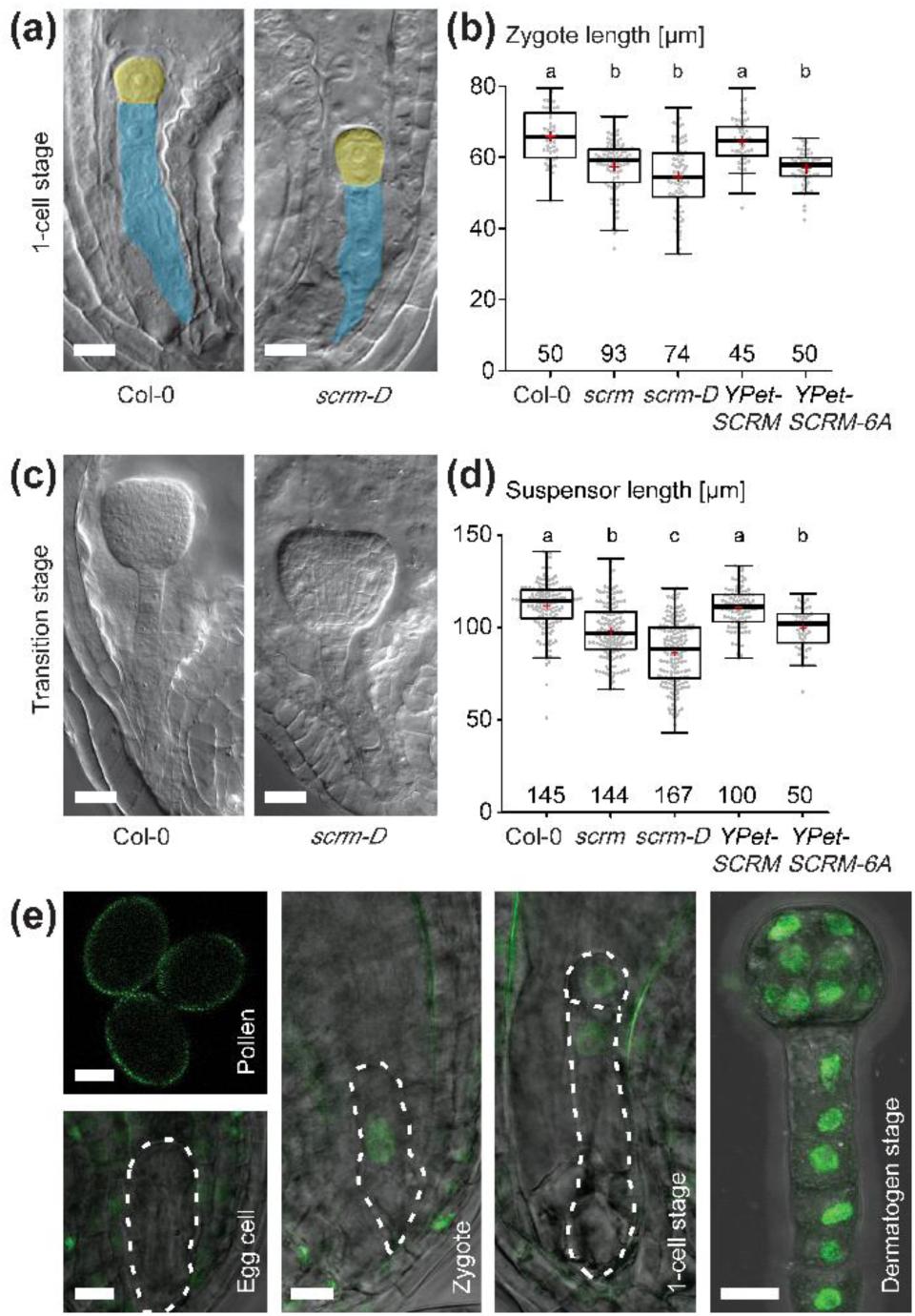
**SCRM regulates zygotic asymmetry and patterning via phosphorylation.** (a) DIC images of one cell-stage embryos (apical cells false-colored in yellow and basal cells colored in blue). The *scrm-D* mutation interferes with MPK6 interaction and inhibits MPK6-dependent phosphorylation. Scale bars, 10 µm. (b) Boxplot diagram of zygote lengths (sum of apical and basal cell lengths at 1-cell stage). (c) DIC images of representative transition stage embryos. Scale bar, 20 µm. (d) Boxplot diagram of suspensor lengths in transition stage embryos. In (b, c), the sample size is given above the x axis. Center lines show the medians; box limits indicate the 25th and 75th percentiles; whiskers extend 1.5 times the interquartile range from the 25th and 75th percentiles; red crosses represent sample means; data points are plotted as gray dots. Letters above boxes refer to individual groups in a one-way ANOVA with a post hoc Tukey test (p < 0.05). (e) Confocal microscopy images of SCRM-YPet in gametes and early embryos. Cell shapes are outlined with dotted line. Scale bars, 10 µm.

The phenotype of the *scrm-D* allele suggests that MPK3/6-dependent phosphorylation is required for SCRM activity in the early embryo, unlike stomata development, where phosphorylation of the SPCH/SCRM heterodimeric complex leads to ubiquitination and proteolysis.

To gain insight into the protein abundance of SCRM during embryogenesis, we genetically complemented the *scrm* mutant with transgenic constructs containing the *SCRM* genomic locus with either an N-terminal or C-terminal translational fusion to the *YPet* fluorescent protein coding sequence. Both constructs complemented the *scrm* mutant phenotypes in the embryo and epidermis (Fig. 3b, d; Fig. S4a, b). Microscopic analysis of the *SCRM-YPet* reporter lines revealed no or very low expression in pollen and egg cells (Fig. 3e). After fertilization, there was an immediate increase in YPet signal in the zygote nucleus, which can also be observed in all cells of the later embryo (Fig. 3e). These findings are consistent with published transcriptomic data from isolated embryos (Fig. S1a) (Zhao *et al*., 2019) and the zygotic and recessive phenotype of the *scrm* mutant.

Observation of YPet fluorescence at the 1-cell stage revealed no obvious difference in the amount of SCRM-YPet protein in the nuclei of the two daughter cells (Fig. 3e). This further supports our observation that the control of SCRM-YPet activity in the early embryo does not rely on cell type-specific degradation and therefore differs from the mode of regulation during stomata development.

The SCRM protein is predicted to contain six potential MAPK phosphorylation sites (Li *et al*., 2017; Zhao *et al*., 2017). To determine whether SCRM function depends on phosphorylation by the YDA pathway, we mutated all six potential phosphorylation sites as previously described by Li et al., 2017 to generate phosphorylation-incompetent (serine or threonine to alanine) SCRM versions and designated them YPet-SCRM 6A.

*YPet-SCRM 6A* was unable to suppress the embryonic defects (Fig. 3b, d), when expressed under the control of the endogenous promoter, consistent with the notion that phosphorylation is required for SCRM function. This finding was corroborated in a previously reported transgenic line carrying a similar phosphorylation-impaired version of SCRM, but with a GFP tag at the C-terminus (Li et al., 2017, ICE1 6A-GFP; Fig. S4c, d). Consistent with our results for the N-terminally YPet-tagged version of SCRM 6A, this previously published line did not complement *scrm* mutant embryo defects (Fig. S4c, d), whereas it resulted in an increase in stomatal index confirming that the construct is functional (Fig. S4a, b).

These results demonstrate that the embryonic function of SCRM is dependent on its interaction with MPK3/6 and MPK3/6-dependent phosphorylation. Notably, the observed gain-of-function stomatal phenotypes and loss-of-function embryonic phenotypes of non-phosphorylated SCRM are consistent with the results obtained with *scrm-D*, highlighting the differential regulation by the ERECTA-YODA pathway in different developmental contexts.

## Discussion

SCRM, also known as ICE1, has been shown to be involved in multiple biological processes in plants, including freezing tolerance, seed dormancy, endosperm breakdown and stomata development (Chinnusamy *et al*., 2003; Kanaoka *et al*., 2008; Denay *et al*., 2014; Hu *et al*., 2019; MacGregor *et al*., 2019). Previous publications have confirmed that it functions as the downstream target of a MAPK pathway (Li *et al*., 2017; Zhao *et al*., 2017), and detailed analysis in stomata development places SCRM downstream of the YDA-MKK4/5-MPK3/6 MAPK cascade, where it cooperates with the transcription factor SPEECHLESS (SPCH) to control the asymmetric entry division and amplifying divisions to initiate meristemoid cells (Kanaoka *et al*., 2008; Putarjunan *et al*., 2019). In zygotes, the ERECTA-YODA pathway phosphorylates WRKY2 to mediate the expression of the basal identity gene *WOX8* (Ueda, Minako *et al*., 2011; Ueda *et al*., 2017). We found that the expression pattern of *SCRM* shows early zygotic activation at the RNA and protein level, consistent with the model that ERECTA-YODA signaling occurs immediately after fertilization (Bayer *et al*., 2009; Neu *et al*., 2019). The observed similarity of defects in zygotic elongation, zygotic polarity, and suspensor length between *scrm* and *ssp-6* suggests that they both function in the same pathway, a possibility further supported by the stronger phenotypes observed in *scrm ssp-6* mutants that resemble those of the *yda* mutant. Further evidence for this mutual involvement is provided by our finding that SCRM acts as a transcriptional activator of *WOX8*, a known target gene of the embryonic YDA pathway.

Our identification of SCRM as a novel target of the ERECTA-YODA pathway during embryogenesis highlights the multiple roles this protein fulfills in different developmental and stress response pathways (Chinnusamy *et al*., 2003; Kanaoka *et al*., 2008; Denay *et al*., 2014; Hu *et al*., 2019; MacGregor *et al*., 2019). Yet, SCRM appears to be universally controlled by MPK3/6-dependent phosphorylation in these different contexts. Thus, SCRM appears to be a central hub for connecting MAP kinase pathways to bHLH transcription factor complexes, as reported earlier for the SPCH-SCRM complex (Lampard, G. R. *et al*., 2008; Lampard *et al*., 2009; Putarjunan *et al*., 2019).

### Phosphorylation-dependent activation of SCRM in zygotes

In the stomatal lineage, MPK3/6 interact with and phosphorylate SPCH and SCRM to reduce their protein stability and transcriptional activity (Lampard *et al*., 2014; Putarjunan *et al*., 2019). SCRM acts as a_scaffold to recruit MPK3/6 via the MAPK docking site and the KRAAM (KiDoK) motif to bridge the MAPKs and SPCH. Underscoring this, a *scrm-D* mutation in the KRAAM motif has been shown to abolish the associations between SCRM/SPCH and MAPKs (Putarjunan *et al*., 2019).

Previous studies have shown that the *scrm-D* loss-of-MPK interaction mutant and the loss-of-phosphorylation transgenic line result in gain-of-function stomata phenotypes due to a lack of phosphorylation-dependent degradation (Kanaoka *et al*., 2008; Li *et al*., 2017; Putarjunan *et al*., 2019). In cold response, the MPK3/6 cascade also acts as a negative regulatory loop to maintain a dynamic SCRM protein level (Li *et al*., 2017; Zhao *et al*., 2017). However, our finding that zygote polarization defects in the *scrm-D* mutant and phosphorylation-deficient transgenic lines were similar to those of loss-of-function mutants (Fig. 2a-e), suggests a phosphorylation-dependent activation of SCRM by the embryonic YDA pathway. We do not detect differential protein degradation of SCRM in the two daughter cells of the first asymmetric cell division. This further supports the notion that SCRM activity is not regulated on the level of protein abundance and that phosphorylation does not lead to protein degradation in the early embryo. This is reminiscent of a positive influence of MPK3/6-dependent phosphorylation on the activity of the FAMA-SCRM heterodimer during guard cell differentiation (Lampard *et al*., 2009). These results suggest that the effect of MPK phosphorylation on SCRM activity is cell type specific, probably depending on interacting partners, such as F-box proteins. Notably, the rice SCRM ortholog OsICE1 was shown to be phosphorylated by OsMPK3 under freezing stress to prevent its ubiquitination by the E3 ligase OsHOS1, leading to the accumulation of phosphorylated OsICE1 and upregulation of *OsTPP1* gene expression (Zhang *et al*., 2017). This is another example that the effect of MAPK phosphorylation on SCRM activity is context dependent. Furthermore, our results raise the question of whether activation of SCRM-dependent bHLH complexes by MAPK phosphorylation is a more widespread phenomenon that may be overlooked in contexts where a potential activation is followed by proteasomal degradation of the SCRM-bHLH complex.

### Independent TFs synergistically polarize the zygote

Our phenotypic comparisons of mutant combinations suggested that the establishment of zygotic polarity requires a regulatory network of synergistic and parallel acting factors. We found an enhancement of the loss-of-function phenotype in the *scrm wrky2-1* double mutant compared to the single mutants, but at the same time we could not show a physical interaction between SCRM and WRKY2 proteins. This suggests that these proteins might belong to independent TF complexes that synergistically regulate *WOX8* expression. Notably, in contrast to the specific expression of *WRKY2* in suspensor cells (Ueda *et al*., 2017), *SCRM* appears to be ubiquitously expressed in all cells of the early embryo. Therefore, at least at later stages of embryogenesis, there are only partially overlapping expression patterns, suggesting that these factors can at least partially function independently.

The HD-ZIP transcription factors HDG11/12 have been identified as additional activators of *WOX8* expression and fulfill a function in zygote polarization, but the links between SCRM and HDG11/12 still need to be further investigated (Ueda *et al*., 2017).

It is worth noting that HDG2, a HD-ZIP IV family paralog close to HDG11/12, has a stomata defect in loss-of-function mutants similar to that of *scrm* (Peterson *et al*., 2013). It would be interesting to elucidate the links between SCRM and the HD-ZIP IV family in stomata development and embryo patterning. In addition, HDG11 has been shown to affect the expression of *ERECTA*, the receptor kinase gene that acts upstream of the YDA-dependent MAP kinase cascade (Guo *et al*., 2019). Since *hdg11* mutants show a similar maternal effect as *er* mutants during embryogenesis, HDG11 may also have a pronounced effect on the expression of the upstream receptor complex (Ueda *et al*., 2017; Wang *et al*., 2021). This may indicate a complex regulatory network that warrants further investigation. The independent activation of the same target gene by different transcription factors, both controlled by the same MAP kinase pathway, may provide an opportunity to integrate multiple signaling inputs and to provide robustness to the signaling output.

In conclusion, this study shows that phosphorylation by the ERECTA-YODA pathway positively influences the activity of SCRM in the early embryo, which differs significantly from the phosphorylation-dependent degradation model in stomata development. It provides a novel example of differential MAPK regulation of a single target protein in different developmental contexts. Our results reveal a parallel regulation of *WOX* gene expression by SCRM and WRKY2 in the context of the asymmetric zygote division. This not only broadens our understanding of how differential gene expression is established in the early embryo but may also provide a new perspective on bHLH-WRKY regulatory networks in the context of asymmetric cell divisions in general.

## Acknowledgements

We would like to thank Yimin Hu and Franca Schneider (Max Planck Institute for Biology, Tübingen, Germany) for experimental help, the microscopy facility of the Max Planck Institute for Biology, Tübingen and the microscopy facility of the Center for Plant Molecular Biology (ZMBP), Caterina Brancato and Dr. Kenneth Berendzen at the Transformation Facility (ZMBP) for protoplast transformations, Dr. Arvid Herrmann and Prof. Keiko U Torii (University of Texas at Austin, USA) and Prof. Yiting Shi and Prof. Shuhua Yang (China Agricultural University) for providing seeds and plasmids. We also thank the Salk Institute Genomic Analysis Laboratory as well as the Nottingham Arabidopsis Stock Centre (NASC) for providing the sequence-indexed Arabidopsis T-DNA insertion mutants. Research in our groups is supported by the German Research Foundation (Deutsche Forschungsgemeinschaft – DFG) grants La606/13-2, 19-1, 20-1 to T.L, BA3356/3-1, BA3356/4-1, and SFB1101/B12 to M.B., the Chinese Scholarship Council (fellowship No. 201806320131 to Y.M.), and the Max Planck Society.

## Author contributions

H.C. and M.B. designed the study with critical comments and modifications by K.W., Y.M. and R.S. H.C., F.X., T.B., K.W., Y.M., and D.S. performed experiments. H.C., F.X., T.B., K.W., Y.M., D.S., T.L., and M.B. evaluated the data and interpreted the experimental results. H.C. and M.B. wrote the initial manuscript draft, all authors commented on and edited the manuscript.

## Data availability

Data of this study is deposited in the Supporting Information of this article and available upon request from the authors.

