## Supplementary Figures and Tables for "Phosphorylation-Dependent Activation of the bHLH Transcription Factor ICE1/SCRM Promotes Polarization of the Arabidopsis Zygote"

### Supplementary information

### Supplementary figures

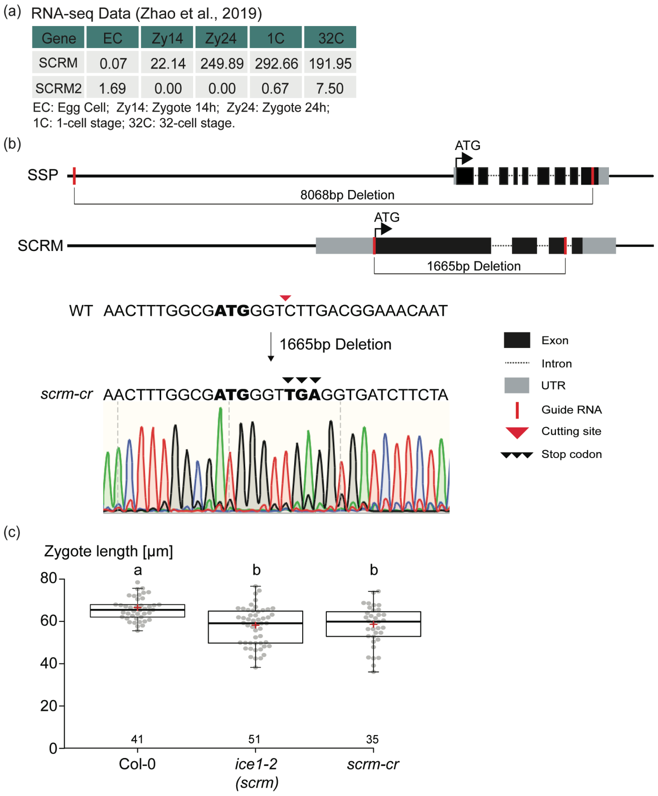


**Figure S1:** Embryonic expression profiles and mutant alleles.

(a) Publicly available RNA-seq expression profiles in the early embryo for *SCRM* and *SCRM2* as reported in Zhao *et al*., 2019. (b) Schematic depiction of CRISPR/Cas9-mediated *ssp-6* and *scrm-cr* mutant alleles. (c) Boxplot diagram of zygote lengths. The sample size is given above the x axis, genotype below. Center lines show the medians; box limits indicate the 25th and 75th percentiles; whiskers extend 1.5 times the interquartile range from the 25th and 75th percentiles; red crosses represent sample means; data points are plotted as gray dots. Letters above boxes refer to individual groups in a one-way ANOVA with a post hoc Tukey test ($p < 0.05$).

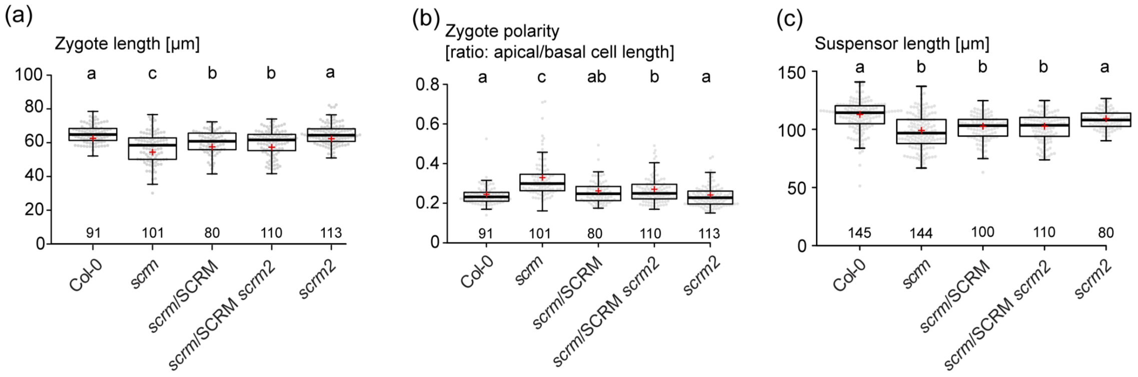


**Figure S2:** Phenotypic analysis of *scrm* and *scrm2* mutant combinations.

(a-c) Boxplot diagram of zygote lengths (a), zygote polarity (b) and suspensor length at transition stage (c). The sample size is given above the x axis, the genotype is given below. Center lines show the medians; box limits indicate the 25th and 75th percentiles; whiskers extend 1.5 times the interquartile range from the 25th and 75th percentiles; red crosses represent sample means; data points are plotted as gray dots. Letters above boxes refer to individual groups in a one-way ANOVA with a post hoc Tukey test ($p < 0.05$).

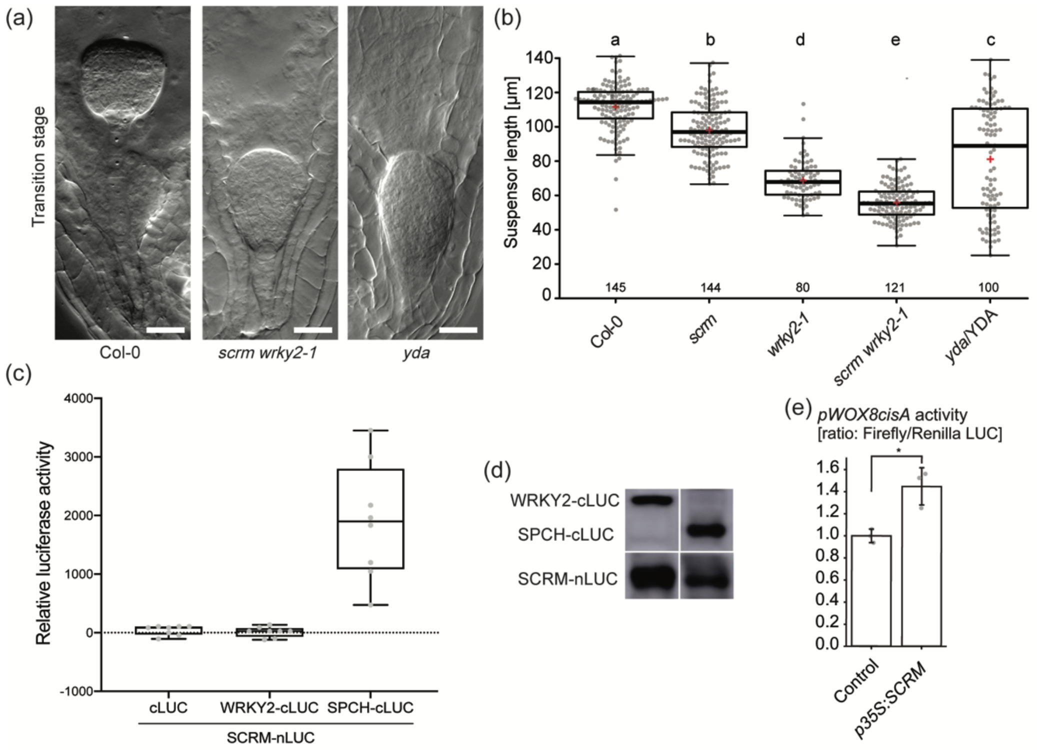


**Figure S3:** SCRM and WRKY2 act synergistically in promoting *WOX8* expression.

(a) DIC microscopy images of transition stage embryos. Genotypes are given below figure panels. Scale bars, 20µm. (b) Boxplot diagram of suspensor lengths at transition stage. The sample size is given above the x axis, the genotype is given below. Center lines show the medians; box limits indicate the 25th and 75th percentiles; whiskers extend 1.5 times the interquartile range from the 25th and 75th percentiles; red crosses represent sample means; data points are plotted as gray dots. Letters above boxes refer to individual groups in a one-way ANOVA with a post hoc Tukey test ($p < 0.05$). (c) Split-Luciferase assay to test protein interaction. Boxplots of 8 replicates for each protein combination. Center lines show the medians; box limits indicate the 25th and 75th percentiles; whiskers extend 1.5 times the interquartile range from the 25th and 75th percentiles; data points are plotted as gray dots. Protein expression was confirmed by Western Blot (d). (e) Dual luciferase activity assay in protoplasts of suspension cell cultures. Data is illustrated as means ± SD of three independent experimental replicates, asterisk indicates statistical differences in Student's t-test ($p<0.05$).

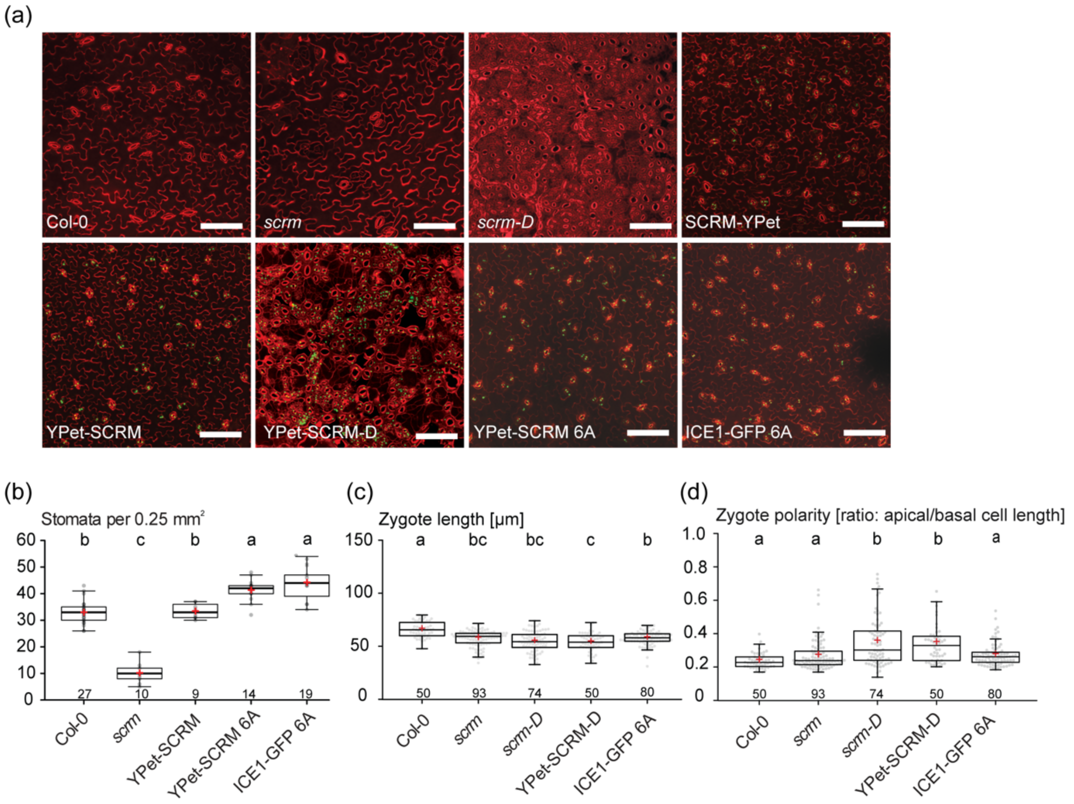


**Figure S4:** Stomatal and embryonic phenotypes in transgenic SCRM variants

(a) Confocal images of abaxial cotyledon of seedlings 5 days post germination (5dpg). Genotypes are given in the figure panels. Scale bar, 100 µm. (b-d) Boxplot diagram of guard cell density in SCRM variants driven by native promoter (numbers of guard cells are measured in 0.25mm$^2$ regions; in b), zygote lengths (c) and zygote polarity (d). The sample size is given above the x axis, the genotype is given below. Center lines show the medians; box limits indicate the 25th and 75th percentiles; whiskers extend 1.5 times the interquartile range from the 25th and 75th percentiles; red crosses represent sample means; data points are plotted as gray dots. Letters above boxes refer to individual groups in a one-way ANOVA with a post hoc Tukey test ($p < 0.05$).

**Supplementary table1**

Genotyping primer list

| Primer Name | Sequences |
|---|---|
| scrm(SALK_003155)-LP | ATTCTTTGCTCTGCCTCTTCC |
| scrm(SALK_003155)-RP | TTTGTAGGGCCTTTGTTGTTG |
| wrky2-1(SALK_020399)-LP | CCAAGAATTTGGCTGAATCTC |
| wrky2-1(SALK_020399)-RP | TGTTAGAACACGAATCACCCC |
| scrm2-1(SAIL_808_B10)-LP | TGCATCCGACATCTTTTATCC |
| scrm2-1(SAIL_808_B10)-RP | TTGTTGTTGTTGCTGTGAAGC |
| yda-11(SALKseq_078777)-LP | GGGTTGTTGATTAGTAAACCATAT |
| yda-11(SALKseq_078777)-RP | TAGTAGGAGACCCAGTAGTT |
| LBb1.3 | ATTTTGCCGATTTCGGAAC |
| SAIL-LB3 | TAGCATCTGAATTTCATAACCAATCTCGATACAC |
| scrm-cr-LP | TATATAAAGACAGAGACAGTGACAACAGAGAC |
| scrm-cr-MRP | AGTAGGCTGTGCACCAGACGATGCT |
| scrm-cr-RP | GATCATACCAGCATACCCTGCTGTATC |
| ssp-6-LP | ATGGTTCTGGTTGGTCCAGGGGA |
| ssp-6-MLP | TGTTCGAAGAAACGGTGATCGAGC |
| ssp-6-RP | ACTGGTCTTGGCAAAAGGGAGATTCA |

**Supplementary Table2**

Cloning and mutagenesis primer list

| Primer Name | Sequences |
|---|---|
| SCRM-YPet-InF | TAGAACTAGTGGATCCAGACAACCGGACCACCGTCAATAACAT |
| SCRM-YPet-InR | CACCATGGATCCGCCTCCACCGATCATACCAGCAT |
| pSCRM(5134)-InF | ATGATTACGAGCTCTCTAGAACTAGTCCTAGGATCTAAAGCCACCT CATTTGC |
| pSCRM(5134)-InR | CAACTCCTCACCTTTAGACACCATCGCCAAAGTTGACACCTTTACC |
| YPet-SCRM-InF | ATGAACGAGTTGTATAAAGGGCCCATGGGTCTTGACGGAAACAATG GTGG |
| YPet-SCRM-InR | GCCAAGCTGACGTCAGGTACCTAACCGCCATTAACTATGTCTCCTC |
| SCRM-D-muF | [Phos]-TTCCAGAAACATGCAGCTATGCGTCAGAGCTCT |
| SCRM-D-muR | [Phos]-GCATAGCTGCATGTTTCTGGAACAGAGTAGGCT |
| SCRM-S94A-muF | [Phos]-TCTTCTTCTTGTGCTCCTTCTCAAGCTT |
| SCRM-S94A-muR | AAGCTTGAGAAGGAGCACAAGAAGAAGA |
| SCRM-S203A-muF | [Phos]-GGAAGGTTTTGGTGCTCCTGCTAATGGT |
| SCRM-S203A-muR | ACCATTAGCAGGAGCACCAAAACCTTCC |
| SCRM-T366A-muF | [Phos]-TGAACTTGAGTCAGCTCCTCCTGGATCT |
| SCRM-T366A-muR | AGATCCAGGAGGAGCTGACTCAAGTTCA |
| SCRM-T382A T384A-muF | [Phos]-AAGCTTCCATCCGTTGGCACCTGCACCGCAAAC |
| SCRM-T382A T384A-muR | GTTTGCGGTGCAGGTGCCAACGGATGGAAGCTT |

| | |
|---|---|
| SCRM-S403A-muF | [Phos]-GTTGTGTCCCTCTTCTTTACCAGCTCCTAAAGGCCAGCAA |
| SCRM-S403A-muR | TTGCTGGCCTTTAGGAGCTGGTAAAGAAGAGGGACACAAC |
| SCRM-Nluc-InF | GGGACGAGCTCGGTACCATGGGTCTTGACGGAAAC |
| SCRM-Nluc-InR | GTACGAGATCTGGTCGACGATCATACCAGCATACC |
| WRKY2-CCluc-InF | CGGGGGACGAGCTCGGTACCATGGCTGGTTTTGATGAAAATGTTGC |
| WRKY2-CCluc-InR | ACGAGATCTGGTCGACAATCTGAGGTAATCTACTCATGATCTGGTTATATACC |
| SPCH-CCluc-InF | CGGGGGACGAGCTCGGTACCATGCAGGAGATAATACCGGATTTTCTTG |
| SPCH-CCluc-InR | ACGAGATCTGGTCGACGCAGAATGTTTGCTGAATTTGTTGAGC |
| SCRM-JIT60-InF | CTTGGCTGCAGGTCGACGGATCCATGGGTCTTGACGGAAACAATG |
| SCRM-JIT60-InR | ATTTCAGCGTACCGAATTCTCAGATCATACCAGCATACCCTGCT |
| bHLH35-JIT60-InF | TTGGCTGCAGGTCGACATGGAGGATATCGTCGACC |
| bHLH35-pJIT60-InR | TCAGCGTACCGAATTCTTAGTAAAGAGAGTCGATG |
| cisA-LUC-InF | AATTCCTGCAGGGATCCATAGTCAAATTAGATTAT |
| cisA-LUC-InR | GAAGGGTCTTGCACTAGTAATTCCTACTCTTAAAT |
| 35Smini-LUC-InF | CCGGGGGATCCACTAGTGCAAGACCCTTCCTCTAT |
| 35Smini-LUC-InR | TTTGGCGTCTTCCATGGGTCGTCCTCTCCAAATGA |
| pWOX8-LUC-InF | CGGGGGATCCACTAGTCATTTCCTTGCAAAAACTC |
| pWOX8-LUC-InR | TTGGCGTCTTCCATGGGATGATGGTGTAATGATGATAATCGAGAGCT |
| ICE1-Nluc-InF | GGGACGAGCTCGGTACCATGGGTCTTGACGGAAAC |

| | |
|---|---|
| ICE1-Nluc-InR | GTACGAGATCTGGTCGACGATCATACCAGCATACC |
| WRKY2-CCluc-InF | CGGGGGACGAGCTCGGTACCATGGCTGGTTTTGATGAAAATGTTGC |
| WRKY2-CCluc-InR | ACGAGATCTGGTCGACAATCTGAGGTAATCTACTCATGATCTGGTTATATACC |
| SPCH-CCluc-InF | CGGGGGACGAGCTCGGTACCATGCAGGAGATAATACCGGATTTTCTTG |
| SPCH-CCluc-InR | ACGAGATCTGGTCGACGCAGAATGTTTGCTGAATTTGTTGAGC |
